# Cerebral Organization in the trilobite *Yunnanocephalus yunnanensis*

**DOI:** 10.64898/2026.10.22.761110

**Authors:** Nicholas J. Strausfeld, Frank Hirth

## Abstract

The clade Trilobita is recognized for its extraordinary morphological diversifications amidst a highly constrained body plan. This raises the question of whether trilobite nervous systems were conservative or comparably diverse. However, despite the identification of thousands of trilobite species and increasing evidence for neural preservation in other Paleozoic euarthropods, the recognition of a fossilized trilobite brain has remained elusive. Here we describe the identification of a trilobite brain and its spatially constrained position within its cranidium. An unusually preserved specimen of the lower Cambrian trilobite *Yunnanocephalus yunnanensis* reveals a bilaterally organized organ connected to its lateral eyes. Comparisons to the brain of a trilobitomorph xandarellid support the interpretation of this organ as a modest cerebrum comprising prosocerebral, protocerebral, and deutocerebral domains, an organization typifying the euarthropod cerebral ground pattern. These findings suggest that the cerebrum of *Yunnanocephalus* could have supported the multimodal sensory processing and behavioral repertoires of trilobites, while retaining the deep evolutionary conservation of brain organization across euarthropod lineages.

## Introduction

Although trilobites have diversified into approximately 20,000 morphologically documented species (Adrian, 2011), their essential biological blueprint, or ground pattern, has remained highly constrained. For example, albeit rarely fossilized, the paired appendages of a trilobite are notable for having the same morphology at every true segment of the cephalon and every true segment of the thorax: thus, homonomous organizations. By contrast, species belonging to the two other major euarthropod clades, chelicerates and mandibulates, are notable for their appendicular divergences. The leitmotiv of an invariant ground pattern is important when pondering how diverse trilobite nervous systems might have become or how simple they might have remained through geological time. And because the brain we describe here is very modest, one wonders whether it represents the full extent of trilobite cerebral organization. Might the more flamboyantly decorated species – and there were very many variations – have possessed greater cerebral complexity? Might trilobite eyes – in some species enormous, and in others optically bizarre – have served similar or uniquely distinct protocerebra? Might the brains of potentially swimming predators have differed from those of their relatives that foraged on the seafloor?

Trilobites have held an allure since the stone age. Paleoanthropologists have found evidence of trilobites’ societal worth across continents (Fernández-Fernández et al., 2025). Archeological evidence suggests they were collected as desirable objects and used as ornaments, probably also as talismans (St. John, 2007). Yet the broad appeal of fossils as objects of study emerged only in the early eighteenth century, a time of growing public wonder of scientific discovery and an appreciation of the natural world (Holmes, 2010). Natural historians and collectors in the early eighteenth century variously claimed fossils to be the shells of bivalve molluscs, limpets, or even the preserved elytra of beetles, which was Linnaeus’ assessment of what we recognize today as trilobites (St. John, 2007). But it was a theologian who in 1771 gave this unique group of fossils not only their name but also notional behavioral attributes

Johann Ernst Immanuel Walch (1725-1778) might be considered a predecessor of a movement that developed in late eighteenth century Germany – an episode of romantic natural science (*romantische Naturforschung*) during which scientific discoveries were inseparable from philosophical and poetic interpretation. Walch spent his intellectual life at the University of Jena, which later would become the intellectual beacon of romantic science (Richards, 2002). He developed his reputation there as a polymath versed in theology, mathematics, linguistics, and the natural sciences, amongst other disciplines, becoming an official professor of many of them. Intrigued by fossils and having viewed a famous collection of ‘antiquities’ when visiting Florence during his ‘Grand European Tour’ (Molitor, 2020), Walch established his own collection and was in contact with other collectors. A fortuitous gift of a large collection of copper engravings of fossils and minerals (Kihm and St John, 2007), motivated him to compile a four-volume treatise *Die Naturgeschichte der Versteinerung* (“The Natural History of Petrifactions”). This work, translated into English in 2007 by Robert Kihm and the paleontologist James St. John, informs us of Walch’s percipient recognition of morphological features that inspired his denomination ‘Trilobite’ to fossilized creatures that shared conspicuously defining attributes: a head “*greatly resembling that of a spider, its back divided into three lobes and garnished with testaceous rings much like the tail of a crayfish, and with a large tail extremity equally divided into three lobes.*” (Walch 1771). Kihm and St. John’s translation reveal the depth of Walch’s thoughts about the natural significance of trilobites in their marine habitat. For example, he proposed that by enrolling its body a trilobite could defend itself. He ascertained that paired structures rising from the head, previously thought to be horns, were a combination of lateral ridges and paired eyes. Walch suggested that different trilobites were distinguished morphologically regarding their patterning of features of the head, trunk, and pygidium – morphological traits utilized today for cladistic analyses of trilobite relationships.

Although Walch was certain that there were no trilobites alive contemporaneously, his comments about trilobite mobility and actions indicate an acute observer of living arthropods. He suggested trilobite’s similarity to living isopods not only because they could vulvate (enroll), but also because they shared overall morphological simplicity. As his translators point out, Walch’s assertion that trilobites were extinct was astonishing coming from a serious theologian decades before the intellectual reality of extinction was accepted. Doubts even persisted after a book published in 1859 revealed its author’s puzzlement that: “*it cannot be doubted that all the Cambrian and Silurian trilobites are descended from some one crustacean, which must have lived long before the Cambrian age, and which probably differed greatly from any known animal*” (Darwin, 1859). If Darwin’s perplexity persists, it is not because trilobite evolution signalled a steady-paced rather than an explosive early Cambrian (Paterson et al., 2019); or that phylogenetics and paleontology identify mud-dwelling vermiform predators equipped with mouths typifying lobopodians as the most likely stem ecdysozoans (Howard et al., 2020; 2022). Rather, it is the challenge of being able to align trilobites with all the other stem euarthropods such that a pan-euarthropod ground plan emerges.

## Results

### Diversity within a conserved ground pattern

As emphasized by Richard Fortey (2000), trilobites exemplify extraordinary morphological diversification while retaining a highly conserved ground pattern over nearly 300 million years of evolution. This differs markedly from that pertaining to many other Cambrian euarthropods whose fossils do not easily allow their designation as ancestral to any specific order of crown mandibulates represented today.

It is fair to say that irrespective of trilobite diversification, involving peculiarly shaped pygidia, exaggerated features of the cranidium, elaborate spines, or grotesque wrought-iron-like excrescences, no fossil radically departs from the ground pattern to the extent that one cannot be sure it is a trilobite (Fortey, 2000; Fortey and Whittington, 1989). The frontal and lateral borders of the cephalon might be smooth in one species, whereas the next fossil encountered may have the border extending far forwards where ruffles and perforations are interpreted as a sieve for a filter-feeding existence (Fortey and Owens, 1999). As weird as such structural variations might be, the identity of the fossil is unmistakably trilobitic. A dome-like cephalon enormously enlarged above a swollen underside, crop and muscle scars on the inside of the glabella’s walls suggest a habitual predator (Lerosey-Aubril and Peel, 2018), whereas unmistakably docile species have evolved signature deterrents: armor from which extend fantastic hooks, prongs, sturdy spikes and trident-like shovels (Fortey, 2004). No matter how extravagantly different, those fossils are instantaneously identified as a mere modification of the body plan. In short, however elaborate or reduced a fossil may appear, it will never be mistaken for anything other than a trilobite if it is one.

### Sensory systems and behavioral capacity: visual diversity

Richard Fortey in his millennial book describes his fascination with trilobite eyes. This preoccupation is understandable because the eyes of some species comprise huge screen-like ensembles of tiny hexagonal facets whose combined fields of view would be panoramic. In swimming species paired holochroal eyes can be cyclopean (Fortey, 1985), fused at the midline, as are the compound eyes of extant Odonata (dragonflies). Faceted eyes ranged from conventional forms to the remarkable schizochroal eyes, a radical modification of the ommatidial design exclusive to the members of Phacopina. Schizochroal eyes have attracted their own literature (Levi-Seti, 1993; Clarkson et al., 2006) not least because the same optical organization has evolved convergently in another order of arthropods, the parasitic insects called Strepsiptera (Buschbeck et al., 2003; Buschbeck, 2005). If the optics of the strepsipteran eye equates to that of the trilobite schizochroal eye, then an eye comprising 100 lenses could view a hundred identical views, yet its underlying neural network encode only motion direction using information of the spaced photoreceptors clustered beneath each individual lens (Maksimovic et al., 2007).

### Mechanosensation, chemosensation, and proprioception

The eyes of trilobites have been used as notional windows through which the observer can perceive a dynamic view of the trilobites’ world (Fortey, 2000). Yet we really do not know how accurately trilobites reconstructed their visual world-its contrasts, colors and motion-if indeed they did at all. Nor would trilobite eyes have been the sole ‘witness of evolution’. Some eyeless phacopine species from abyssal environments imply that other sensory modalities must have also been in play. High-power microscopy has revealed that despite its biomineralized calcite exterior, the trilobite exoskeleton had microscopic slits and wart-like hillocks, suggesting thousands of strategically positioned sensilla (Hammer, 2000; Pillola, 1996; Wilmot, 1991; Miller. 1976). As in extant marine arthropods, each sensillum would have been populated by multiple sensory neurons sensitive to touch, pressure, and even salinity if a species’ habitat was tidal. Sensilla dispersed strategically across the tergites ensheathing the cephalon, thorax and pygidium would have provided vital information about the trilobite’s immediate surroundings. Trilobites must have been highly sensitive to tactile and chemosensory stimuli. Paired flagelliform antennules, comprising scores of articles, extended forwards from the cephalon and, in some species, were at least as long as the body. The only exception (thus far) is a species whose short, clavate antennules bore rows of regularly placed, dome-shaped features, comparable to insect campaniform sensilla that act as proprioceptive receptors (Edgecombe and Fortey, 2022).

Comparisons with extant species of eucrustaceans may give clues about what trilobites could perform behaviorally. For example, the caudal fused segments comprising the pygidium were in some supposedly reef-dwelling trilobites modified as huge fan-shape structures (Fortey, 2000; Kobayashi and Hamada, 1965; Ludvigsen, 1987). These may have been analogues of the telson of living decapod crustaceans. One broadened trilobite ‘tail’ bears tiny hillocks that may represent a field of sensilla capable of triggering pygidial-flip escape responses, as achieved by some crustacean telsons in response to sudden touch or far-field disturbance. Although trilobite biramous appendages were not calcified and are therefore rarely fossilized, preserved examples bear abundant sensilla, comparable to those on the walking legs of extant marine crustaceans.

### Evidence for cerebral preservation in the fossil record

Trilobites did not survive the Permian’s terminal event (Erwin, 1994, Knoll et al., 1996). There are no living relatives. Other members of Artiopoda, the clade to which trilobites belong, had died out long before the Permian. This geological history elicits many questions. How did Trilobita survive so long? How might they have reacted to the sensory world around them? Did they respond to visual motion or even shapes and colors? Did their antennules suggest differences across species in responding to odors or tastes? Could a browsing trilobite have possessed sensory-motor circuits that enabled exploration and the ability to return to a familiar location? If we knew something about their brains, even to discover whether they were capacious or minute, this might further suggest what sensory modalities were represented within different cerebral arrangements and whether such arrangements reflected different biological habitats.

Interpretations that enliven fossilized nervous systems (Fig. 1A-C, E) are usually based on more than single specimens; for example, divergence of brain organization in Cambrian great appendage (GA) arthropods is indicated from nine well-preserved specimens representing three distinct taxa. Neural cladistics of specimens of *Jianfengia multisegmentalis* from the Chengjiang biota (Fig. 1E) suggests it is a proto-mandibulate that had not yet evolved mandibulate gnathal attributes, yet its brain clearly corresponds to the cerebral organization of extant decapods (Strausfeld et al. 2025a). Six specimens belonging to the megacheirans, *Alalcomenaeus* or *Leanchoilia*, are resolved as two kinds of stem chelicerates (Fig. 1B, C). Both feature two pairs of single-lens eyes serving cerebral arrangements resembling those of living juvenile xiphosurans (Tanaka et al., 2013, Lan et al., 2021). Specimens of *Fuxianhuia protensa*, a species assigned to Mandibulata (Aria et al., 2021), were also collected from Chengjiang sites in southwest Yunnan. Their preserved brains confirmed the initial 2012 findings (Ma et al., 2012) that their compound eyes supplied an elaborate optic lobe in which chiasmal axons remained resolvable even after being entombed for half a billion years (Ma et al., 2015). The absence of a second pair of antennae in *Fuxianhuia* (Fig. 1F) is also telling in that this evolved loss of the paired appendages belonging to the first true segment typifies extant myriapods and hexapods. Interpretive reconstructions from such fossil material (Fig. 1) also include a single chelicerate whose cerebral organization indicates it is a marine arachnid, typified by the characteristic backfolding and semi-reversal of the linear arrangement of the three cerebral domains (Fig. 1D; Strausfeld et al., 2025b). Also joining this menagerie is a member of the artiopodan clade Xandarellida with its reconstructed brain and ventral nerve cord (Strausfeld et al., 2026a; Fig. 1A). In the context of understanding early brain diversification in the Cambrian, the occurrence of fossil cerebra providing meaningful data is still extraordinarily rare. Vanishingly rare is that only one of the 20,000 trilobite species has yielded a satisfyingly dark profile suggesting a fossilized brain (Losso et al., 2026b).

**Fig. 1.**
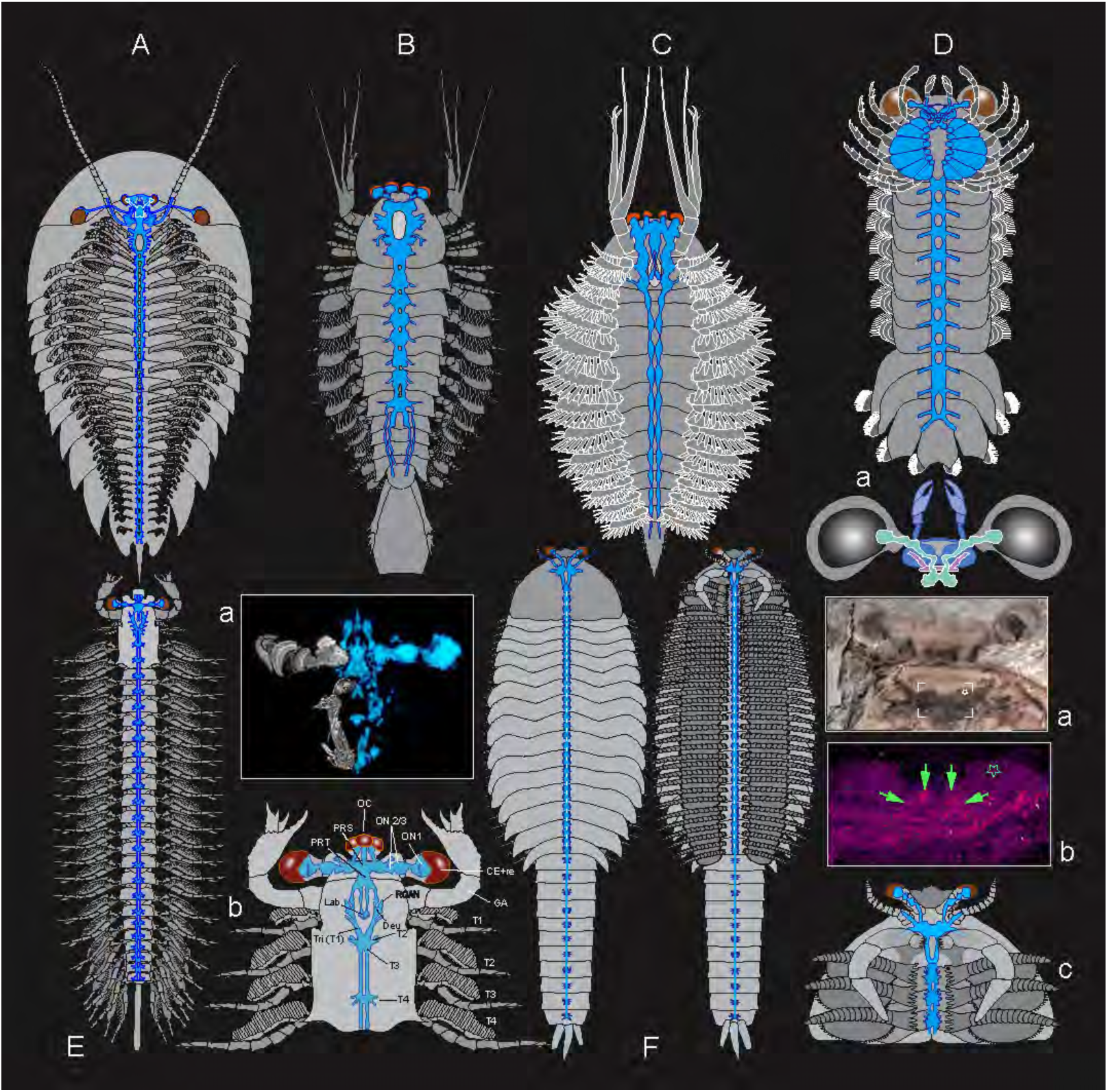
Brains and ventral ganglionated nervous systems of Cambrian Euarthropoda. (Scales have been omitted). **A**. The artiopodan xandarellid central nervous system (CNS, in blue) reconstructed from the fossilized cerebrum of *Xandarella spectaculum* and segmental ganglia from *Cindarella eucalla* (Strausfeld et al., 2026a). **B**. The megacheiran *Alalcomenaeus* and **C** the megacheiran *Leanchoilia* from the Chengjiang biota, both proposed as stem chelicerates (Tanaka et al., 2013; Lan et al., 2022). **D**. The Burgess Shale *Mollisonia symmetrica* initially described as a stem chelicerate. Re-examination of the specimen identifies a specialized cerebrum, shown in **D-a**, where the linear brain organization typifying stem chelicerates is morphogenetically folded over on itself such that the order of the cerebral domains is reversed. The deutocerebral c3 domain that was ancestrally caudal became the most rostral domain (shown blue) with its nerves extending to the paired chelicerae. The ancestral anterior domain (shown green) becomes the most caudal one (Strausfeld et al., 2025b). This organization typifies the morphology of the prosoma and cerebrum of extant arachnids. **E**. Neural characters resolve the megacheiran *Jianfengia multisegmentalis* as a stem mandibulate. **E-a** shows the right side (blue) of the reconstructed fossil brain framed facing the left side (white/grey) of a modern decapod brain. **E-b.** The interpretive drawing of the cerebrum and anterior ventral nerve cord indicates neuropils, nerve branches, and segmental arrangements, all of which support *Jianfengia’s* identity as a stem mandibulate. **F**. The stem mandibulate *Fuxianhuia protensa* shown from above (left) and from beneath (right).). **F-a.** Dark fossil cerebrum (brain) against other pale fossilized structures. **F-b.** An energy-dispersive X-ray spectroscopy image showing the prosocerebrum (green arrows); **F-c**, reconstruction of the brain fused with the second segmental ganglion supplying nerves to the paired sickle-shaped mandibles. The first true segment is not delineated and lacks appendages. An appendage-free first true segment is a defining feature of Hexapoda and Myriapoda referred to as the intercalary segment (Snodgrass, 1952).

### What is preserved inside a trilobite, and what is not

All the examples of fossilized nervous systems represented in Fig. 1 are from euarthopods whose exoskeletons were likely soft, offering taphonomic permeability and thus initial steps in the preservation of soft tissue (Parry et al., 2019; Saleh et al., 2020). Although certain soft tissues are occasionally preserved in some trilobite fossils, these are mostly constrained to ventral appendages, which were not calcified like the body’s tergites (Hou and Hopkins, 2024; Losso et al. 2026a). Internal soft tissue preservation is very rare but has been meticulously documented for the alimentary system: the hypostome, crop, stomach, and other parts leading to the dorsal gut. Preserved parts of the digestive tract have been described (Chatterton et al., 1994, Lerosey-Aubril and Peel, 2018; Lerosey-Aubril et al., 2011, 2012; Kraft et al., 2023; Zhang and Liu, 2026), even to the extent that ingested fragments inform about what the trilobite was consuming before its demise. Some had preyed on soft-bodied animals, whereas others were durophagous or even contained bits of what was likely once a trilobite. Fossilized digestive systems thus provide useful indicators of benthic lifestyles. There is, however, the assumption that the impermeability of the calcite exoskeleton delayed diagenesis and subsequent preservation of tissues, including neural, thought to be more labile than gut. Until now, only one example of soft tissue preservation in a trilobite has been claimed as neural. This is in the early Ordovician trilobite *Anacheirurus adserai* whose cerebral remains are shown as dark rust-red profiles against a mottled pale yellow/grey/blue matrix (Losso et al., 2026b). The specimen shows the first segmental ganglion of the ventral nerve cord attached to a bi-partite area claimed as a cerebrum; this is interrupted by several lacunae one of which is improbably identified as fossilized remnants of an integration center called the ‘central complex’ epitomizing extant stomatopods and pterygote insects, thus asserting Trilobita as crown Mandibulata (Losso et al., 2026b).

### Exceptions to the “dark neural tissue” expectation in fossils

Chemical events underlying fossilization are manifold (Parry et al., 2019; Saleh et al., 2020), and traces of neural tissue are not always resolved as dark. There are two published examples where fossilized nervous tissue appears as a pale deposit against a darker background. One is a fossilized central nervous system belonging to an ancient ‘horseshoe crab,’ a xiphosuran chelicerate. Living horseshoe crabs exist today; they are abundant on the northeast coast of the United States. The fossil species, *Euproops danae,* was retrieved from the late Carboniferous Mazon Creek Lagerstätte in Illinois. Its nervous system had decayed and been replaced with kaolinite, which glistens white against the darker impression of the typical limulid body (Bicknell et al., 2021). The second example is the well-preserved cerebrum and ganglionic ventral nervous system of *Gigamachilis triassicus*, a large Triassic bristletail (Machilidae) from the Monte San Giorgio UNESCO World Heritage site shared by France and Italy (Montagna et al., 2017). In this specimen, pale phosphatized neural tissues are shown against darker structures (for example muscle) backed by a darker matrix. A third example reported here is the fossilized cerebrum of the Lower Cambrian redlichiid trilobite *Yunnanocephalus yunnanensis*.

### The *Yunnanocephalus* specimen and its candidate cerebrum

Science, however, thrives on exceptions. Figure 2 shows an early Cambrian trilobite that provides a third exception to the expectation that fossil neural tissue is usually dark (Ma et al. 2015). *Yunnanocephalus yunnanensis* is a basal Cambrian redlichiid trilobite common amongst the Chengjiang biota. Occasional specimens turn up far from their origins, as did the specimen shown in Fig. 2. Both part and counterpart are nearly transparent. They provide complementary views inside the specimen, that are almost identical but with most material adhering to the part.

**Fig. 2.**
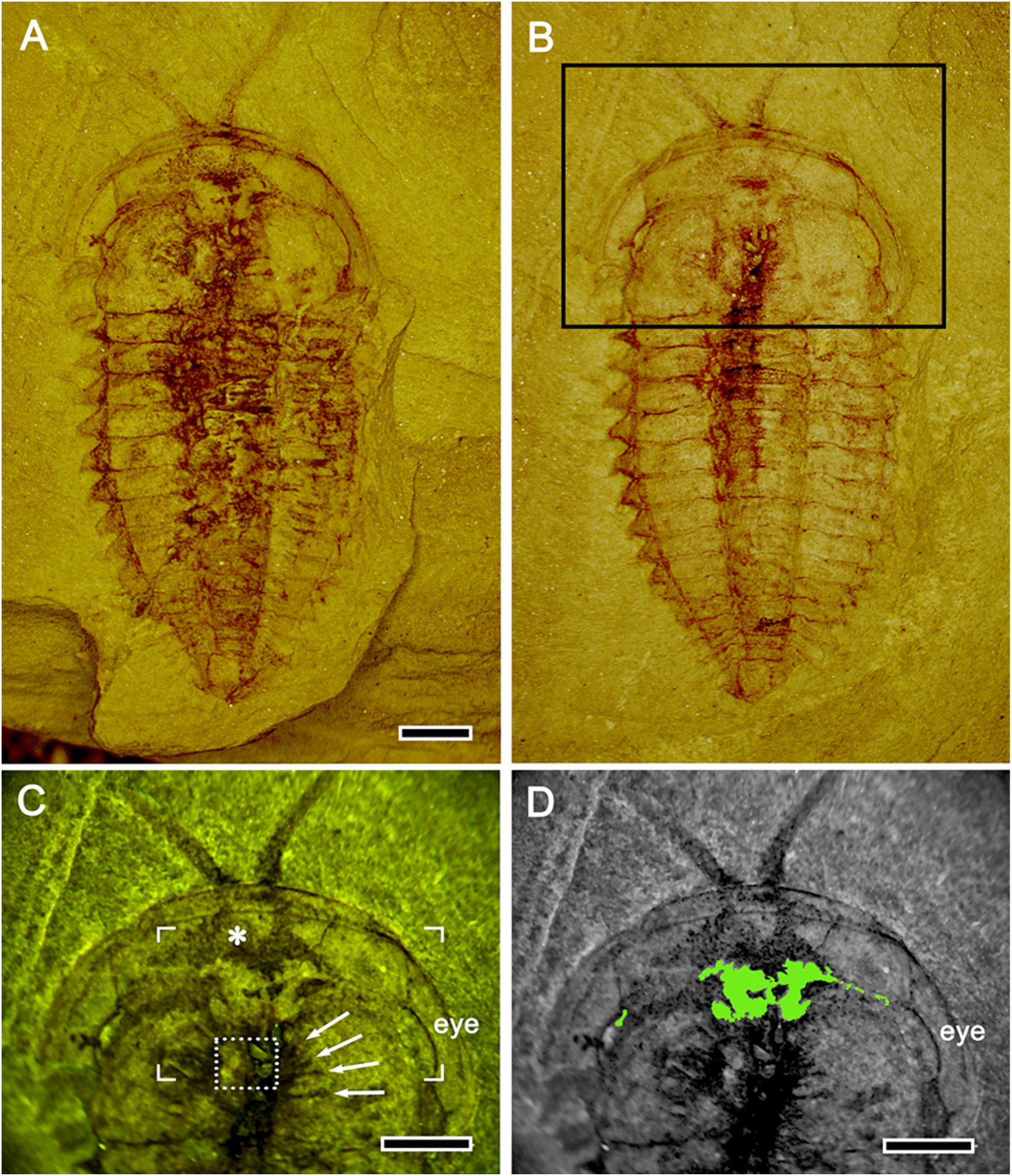
The Cambrian trilobite *Yunnanocephalus yunnanensis.* The part (**A**) and counterpart (**B**) of the Cambrian trilobite *Y. yunnanensis.* In both, the cephalon offers clear resolution of its contents in contrast to the chaotic distribution of fossilized soft material in the thorax. **C**. Enlarged view of the part’s cephalon illuminated by mixed ultraviolet and white light. A pale, near-mirror-symmetric structure overlies a larger dark granulate area (asterisk), also approximating a mirror-symmetric, soft-tissue structure likely to be the underlying hypostome, attached at two locations to the inside of the cuticle. The dotted box at the center denotes ovoid structures that may be associated with the gut. The white arrows indicate traces interpreted as the decayed remnants of the first four post-antennular segmental appendages. The position of the eye is indicated. The corner-bracketed area is the focus of scrutiny depicted in the following two figures. **D.** Green level conversion of the pale structure in panel C. Note the extension leading to the anterior edge of the eye ridge. Scale bars = 2mm.

Figure 2A provides a view into the part, where many components of the thorax, presumably gut and other tissues, have been displaced to provide a compressed muddle. The counterpart in Fig. 2B is a view upward towards the roof of the body. It nicely reveals the three parallel segmented lobes typifying all trilobites first described by Walch (1771). Most of the thorax yields little interpretable information; but the cephalon is different. It speaks volumes. Mixed ultraviolet and white illumination of the part (Fig. 2C) reveal an orderly granular area (indicated by an asterisk in Fig. 2C). To the right, arrows denote depressions indicating the radiating traces of the protopodites of the first four post-antennular appendages. Centered in the image are some ovoid structures that may correspond to glands associated with the gut. The most informative feature is the pair of profiles linked by two commissure-like bridges. The profiles are not quite bilaterally symmetric, and the smaller one on the right extends a branch to its ipsilateral eye. Isolated, by comparison with other fossil cerebra, this feature suggests a bilateral brain. Detailed accounts of alimentary-related skeleton (Lerosey-Aubril et al., 2011; El Albani et al., 2024) and soft tissue fossilization (Eriksson and Terfelt, 2012) provide nothing that would match the pale structure shown in Fig. 2C and isolated in 2D.

### Testing taphonomic distortion

If the left and right profile just described were bilaterally symmetrical in life, they would have represented an organ. Given its position beneath the anterior lobe of the glabella, the most likely candidate in Fig. 2C, D would be a rostral brain comparable to the rostral brains of other Cambrian species (Fig. 1). Closer inspection shows this pale structure nestled over a bilaterally disposed granular area that appears to be attached to the front cuticle at the position where the antennules extend from the head (Fig. 2C). The identity of this granular structure could be best interpreted as the expansive stomach (El Albani et al., 2025). Another interpretation is that it is muscle attached to the base of each antennule thereby enabling active sampling of the surround, comparable to active antennular movements by certain insects (Baba and Comer, 2008; Okada and Toh, 2004).

Could local taphonomic processes, such as displacement under pressure, have produced the observed asymmetries in the cephalon? If the left half of the brain is a distorted version of the right, it should be possible to transform the right half so that it adopts the profile of the left. Figure 3 demonstrates the procedure to test this. We superimposed a rectilinear grid over the right half, which was assumed to be the better-preserved half because of its size and connection to the eye. Both images were flattened into the same plane, and the resultant image (grid plus right half, Fig. 3A) was copied, flipped over the midline, and superimposed on the larger left half (Fig. 3B). This provided a mirror-symmetric construct (Fig. 3B, F). Figure 3C shows the results of using the Photoshop warp function to adjust the grid so that the outline of the right half matches the outline of the left half. Distortions of the grid indicate where expansions likely occurred before diagenesis. Figure 3E shows what the brain would look like if the left half were the better-preserved half. Figure 3E and 3F are two alternative bilateral symmetric profiles, which are proposed here as two versions of a *bona fide* brain. The final test was whether this slightly distorted bilateral structure could confidently be interpreted as a flattened cerebrum?

**Fig. 3.**
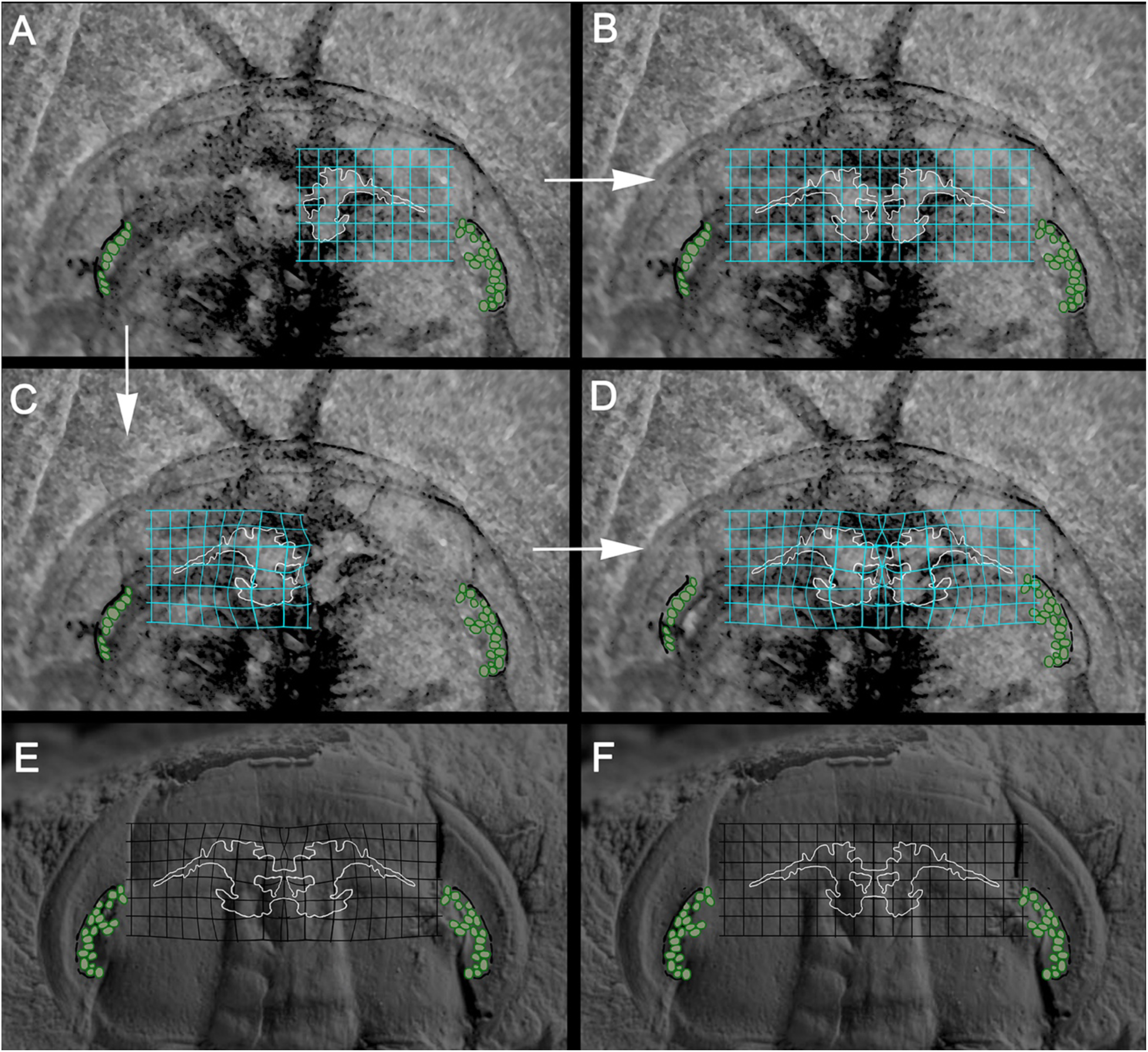
Geometric reconstruction of the cerebrum of *Yunnanocephalus yunnanensis.* **A.** The left half of the cerebrum has a larger footprint than the right half. To determine if the left footprint is due to taphonomic distortion, the right half and its overlying grid was flipped horizontally over the left half in **B** to provide a mirror-symmetrical image approximating the living tissue. **C.** The flipped right half image and its grid were superimposed over the larger left half, and the Photoshop warp functions were used to match the outline of the right half with the outline of the larger footprint. The grid distortion indicates how tissue may have been disproportionately compressed during preservation. In **D** the outline of the left side was flipped horizontally to provide a mirror-symmetrical image. **E, F.** Outlines of the brain, one constructed from the left half (**E**) and one from the right half (**F**), superimposed on the exoskeleton on another specimen of *Yunnanocephalus*, provide some confidence that the putative cerebrum was originally mirror symmetric.

### Comparative validation with *Xandarella*

A specimen of the artiopodan clade Xandarellida, whose lateral-eye organization was described previously (Strausfeld et al., 2016), was recently re-examined with particular attention to dark, mirror-symmetric profiles that reside immediately beneath the dorsal roof of the head shield just anterior to the rostral margin of the hypostome (Strausfeld et al., 2026a). Reconstructed together, serial imaging resolves these fossilized traces as an unambiguous cerebrum associated with the lateral eyes and with paired ocelli situated at the most rostral bridge-like connection between the two brain hemispheres.

Figure 4 enables comparison of the brain of the artiopodan *Xandarella spectaculum* to a simpler but corresponding organization in the cephalon of *Yunnanocephalus yunnanensis*. In Fig. 4A the trilobite brain is shown with one preserved optic nerve (yellow arrows) extending to the lateral compound eye. In the reconstructed brain (Fig. 4B), optic nerves extend to the lateral eyes through the palpebral lobe canal contiguous with the medial eye ridge. In Fig. 4C, the dark fossil profiles of the xandarellid brain exhibit the same arrangement reconstructed in Fig. 4D; in this artiopodan the optic nerves are obscured by the head-shield cuticle within which they project. Thus, in both the xandarellid and the trilobite, the optic nerves are concealed within their respective tergites.

**Fig. 4.**
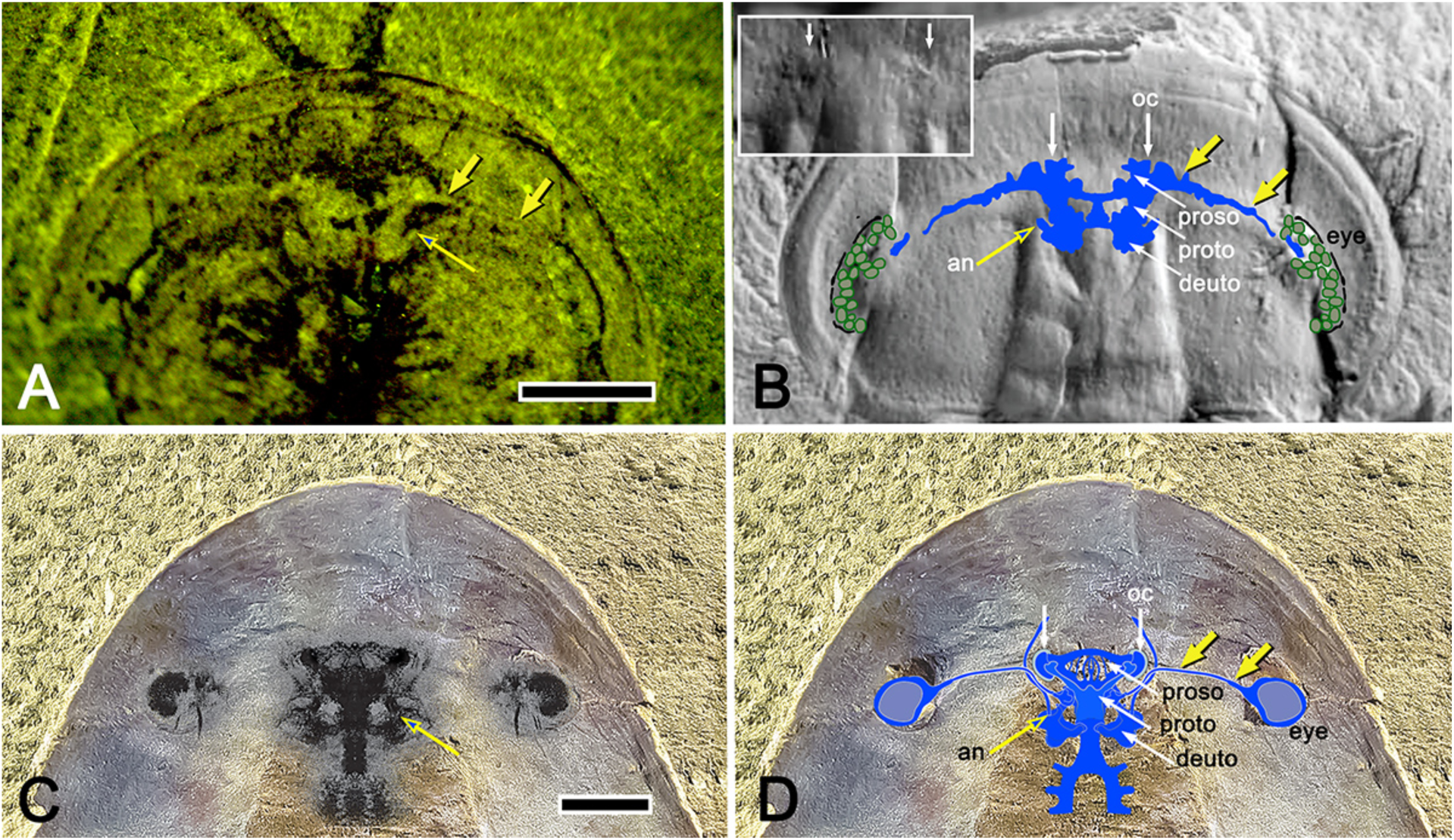
The artiopodan cerebrum in the trilobite *Y. yunnanensis* compared with that of the xandarellid *X. spectaculum*. **A.** The suggested trilobite brain with its optic nerve (yellow arrows) also showing the root of the antennular nerve (open arrow). **B.** The reconstructed bilaterally symmetric cerebrum has been superimposed on another fossil specimen of *Y. yunnanensis* (see Acknowledgements). The inset shows two lenticular dimples suggesting paired ocelli (oc) indicate as two vertical arrows in the main image. Three successive lobe-like divisions of the cerebrum suggest three contiguous domains recognized in extant euarthropods as the morphogenetically defined proso, proto-and deutocerebrum with the root of the antennular nerve (an) indicated. Also indicated are the lateral eyes (eye, green) and their optic nerve (yellow arrows). **C**. The central cerebrum and the flanking lateral eyes of the artiopodan *Xandarella spectaculum* (Strausfeld et al., 2026a) The correspondence of the cerebral and lateral eyes of the xandarellid and trilobite are notable. **D**. The similarities also apply to the location of the paired ocelli (oc), indicated by the vertical white arrows, and to the successive parts of the xandarellid cerebrum. In both the trilobite and xandarellid the connection (yellow arrows) of the lateral eyes to their central neuropils is cryptic. In *Xandarella* the nerves are hidden within a conduit integrated within the overlying tergite; likewise, the trilobite optic nerve extends from its eye to the cerebrum following the course of a homologous conduit (comprising the palpebral lobe and eye ridge). Scale bar for A and B= 2mm; C, D = 5mm.

An exceptionally well-preserved specimen of *Yunnanocephalus*, provided by Prof. Xingliang Zhang (Dai and Zhang, 2008; see acknowledgements), permits precise alignment of the trilobite cerebrum and optic nerve with the crescent-shaped palpebral lobe. In addition, just discernible in the specimen are two ovoid dimples (Fig. 4B, inset), whose positions correspond to the xandarellid paired ocelli. As demonstrated for numerous examples of structures termed ‘frontal’ eyes (Schoenemann and Clarkson, 2023), these structures form part of the ground pattern not only of trilobites, but possibly of Artiopoda as a whole and many other euarthropod lineages. In any event, their presence indicates an important sensory input to the artiopodan prosocerebrum, whereas the lateral eyes identify the protocerebral domain. The presence of paired antennules and their corresponding lobes indicate the third domain of the cerebrum, namely the deutocerebrum.

## Discussion

### Brains align across euarthropods

A comparison (Fig. 5) of the schematized cerebra of the two artiopodans, *Yunnanocephalus* and *Xandarella*, and two mandibulates -the fruit fly *Drosophila* and the aforementioned *Jianfengia multisegmentalis*-illustrates an ancestral morphogenetic organization of the cerebrum that is common to euarthropods, including the folded cerebrum of arachnids (Fig. 1D). This conserved ground pattern of the euarthropod brain comprises three asegmental, genetically determined domains: the prosocerebrum, protocerebrum, and deutocerebrum (Strausfeld et al., 2022, 2026). The labrum and ocellar-type eyes provide sensory input to the prosocerebrum; the compound eyes, to the protocerebrum; and the antennules, to the deutocerebrum. In extant euarthropods the prosocerebrum is populated by distinctive circuits comprising the “central complex,” which supports path integration and spatial memory amongst other directive motor actions. Those circuits are most elaborate in winged hexapods and the predatory stomatopods (Thoen et al., 2017). They are simple in basal pancrustaceans and simplest in myriapods (Strausfeld et al., 2026b). That the fossilized prosocerebrum of *Xandarella* reveals a decussating bridge which is the central complex’s most essential (and basal) circuit, suggests that this organization is likely to be a general feature of artiopodans, including trilobites.

**Fig. 5.**
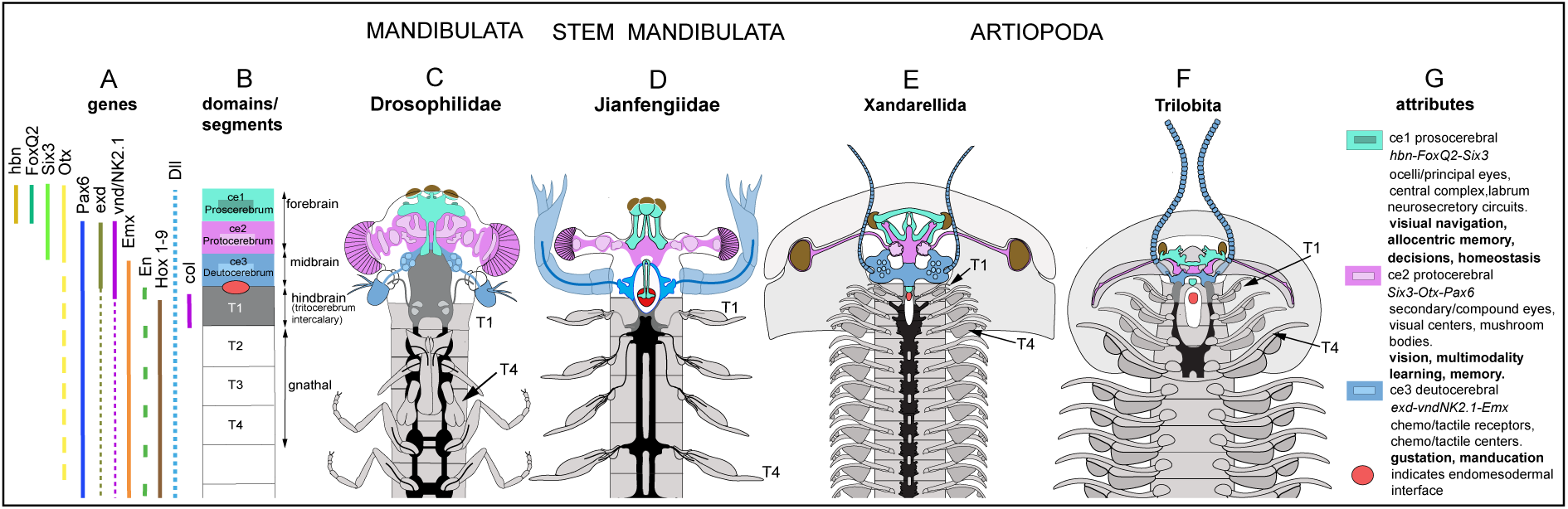
Alignment of extant mandibulate, stem mandibulate and artiopodan brains. **A.** In extant euarthropods combinatorial expression of homologous genes (for source references see Strausfeld et al., 2022) defines three cerebral domains. **B.** These domains are the prosocerebrum (ce1), protocerebrum (ce2), and deutocerebrum (ce3). T1 is the first true segment of the body. The cerebral traits defining each lineage are independent of Hox code-specific determination of segments as T1, and T2-T4, which in extant mandibulates contribute to gnathal appendages. **C-F**. Schematics showing the morphogenetic divisions of the *Drosophila* brain, an extant mandibulate (**C**) aligned with the corresponding neuroanatomical domains of the stem mandibulate *Jianfengia multisegmentalis* (**D**), the artiopodan *Xandarella* (**E**), and the trilobite *Y. yunnanensis* (**F**). **G**. The linear brains of Cambrian and extant euarthropods are characterized by their sensory systems and computational centers. Although anatomists describing trilobites use their own nomenclatures for denoting appendages associated with the cephalon and those associated with the thoracic ganglia, we have used here the standardized terminology that evolved from studies of insect and crustacean embryology. Thus, T1 is the first true post-antennular segment, which in insects is the intercalary segment, but in crustaceans carries the paired (2^nd^) antennae. Segments T2-T4 have greatly diversified across pancrustacea and are involved with feeding, hence called the gnathal segments. In anagrelides and trilobites, the first 4-5 pairs are homonomous segments with biramous appendages.

Further underlining commonalities across euarthropod lineages is that the first four biramous appendages immediately caudal to the cerebrum align across Mandibulata and Artiopoda. *Xandarella* and *Yunnanocephalus* share the basal condition of possessing four homonomous pairs of post-antennular appendages, which in *Yunnanocephalus* (Collantes et al., 2026) originate from beneath the glabellar lobes L3-L0. These correspond to segments T1-T4 of Mandibulata (Fig. 6). The stem mandibulate *J. multisegmentalis* is likewise equipped with homonomous appendages (Fig. 1E), further indicating that homonomous T1-T4 biramous appendages is a basal trait. In crown mandibulates T1-T4 provide divergent appendicular specializations. An example is the paired biramous antennae of segment T1 that denotes eucrustaceans; another is the evolved *absence* of appendages at T1 that hallmarks Myriapoda and Hexapoda, where T1 has been referred to as the “intercalary” segment. Yet across pancrustaceans and chilopods the biramous appendages of segments T2-T4 are specialized for gnathic functions.

**Fig. 6.**
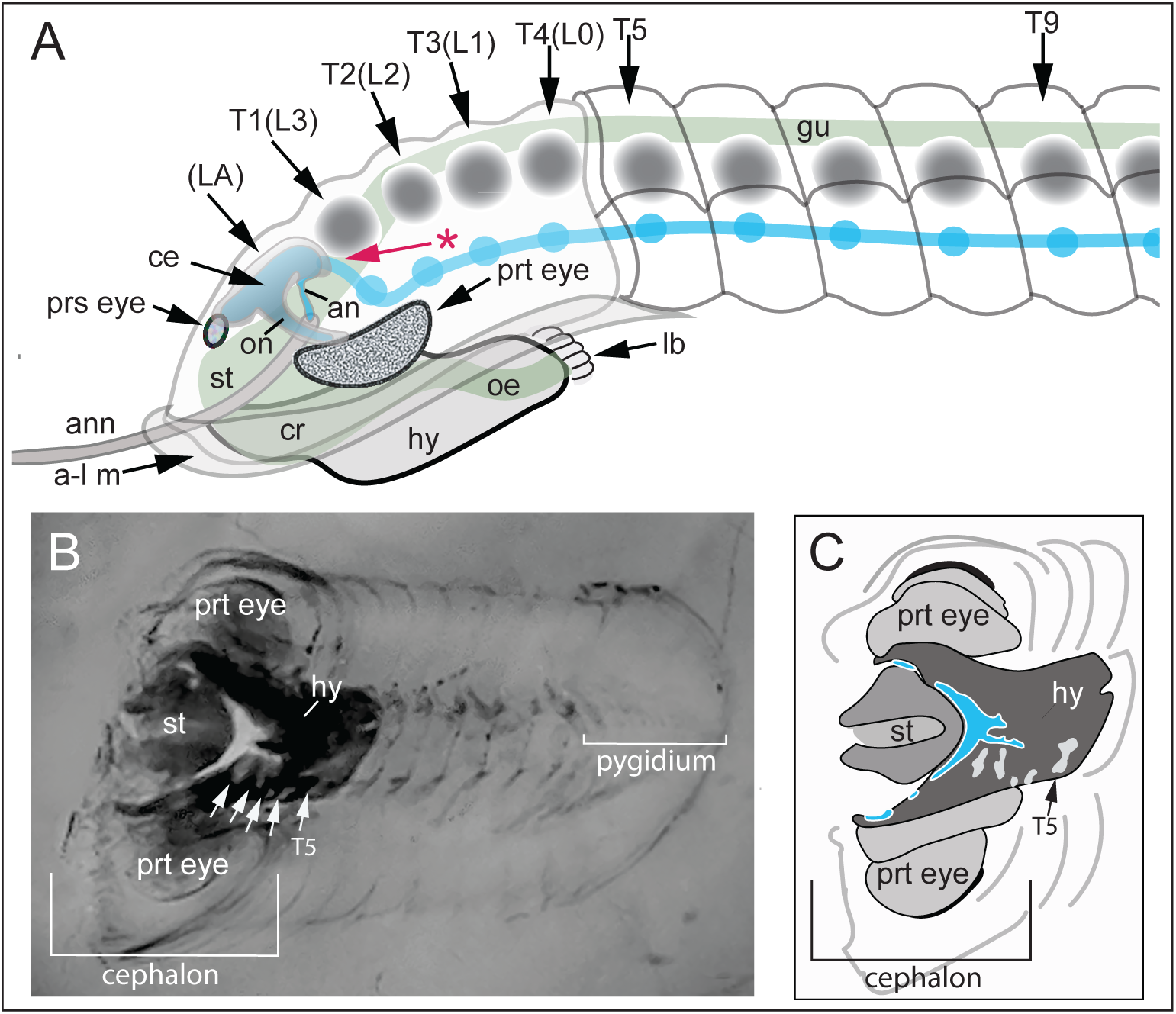
Cerebral and alimentary arrangements within the trilobite cephalon. **A.** Lateral schematic of cephalic elements and their arrangements derived from observations of *Y*. *yunnanensis* (see also, Hou and Hopkins, 2024; Al Albani et al., 2024). The trajectory of the alimentary canal is shown light green; that of the nervous system, light blue. The assumption here is that segmental trilobite ganglia are ventrally disposed as they are across Euarthropoda. The arrangements in *Y*. *yunnanensis* shown in Figs. 2-4 show the brain situated beneath and slightly forwards of the anterior glabellar lobe (LA). It is shown at that location here, above the enlarged hypostome/stomach (st) and crop (cr) of the alimentary canal. The nomenclature L0-L3 used for designating the trilobite glabellar lobes is here shown in parentheses alongside standard abbreviations for true segments, which in this species is T4-T1 forwards. The cerebrum and the first segmental ganglion connect at the level of the endomesodermal interface (asterisked red arrow); thereafter segmental ganglia are ventral. The T1 ganglion is thus the homologue of the tritocerebral ganglion of pancrustaceans. Grey discs indicate the origins of paired biramous appendages (omitted) at segments T1 to T9 (and beyond). Abbreviations: a-l m, anterior-lateral margin; an, antennular nerve root; ann, antennule; ce, cerebrum; cr, crop; hy, hypostome, lb, labrum; oe, esophagus; on, optic nerve; prs eye, prosocerebral ocellus-type eye; prt eye, lateral protocerebral (compound) eye; st, stomach. **B**. Historical X-ray image of the Phacopid *Chotecops* (*Phacops*) *ferdinandi* (modified from Fig. 1g in Schoenemann et al., 2021; see Acknowledgements) provides a top-down view of the cephalic organization described in panel A. Each of the protocerebral schizochroal eyes (prt eye) surmounts a large lateral volume corresponding to a sub-retinal visual neuropil characteristic of Phacopida (Schoenemann et al., 2021). The almost-black volume of the hypostome shows its forward and lateral ‘wings’ flanking the stomach (st). The white crescent-shaped volume and its laterally extending nerves are interpreted as the cerebrum positioned over the hypostome and abutting the rostral stomach, hence disposed at the level of the anterior glabellar lobe. The four segmental glabellar lobes (arrowed) are indicated followed by the T5 segment, thus the first trunk segment. **C.** Main components of the cephalon in panel B rotated by 12° anticlockwise with neural tissue indicated in blue. The length of the specimen is approximately 3 centimeters.

Discussion of the trilobite cerebrum requires consideration of the extensive nomenclature applied to the trilobite exoskeleton (Whittington and Kelly, 1997), which can complicate assessments of artiopodan-mandibulate homologies. A straightforward example is the term ‘antenna,’ potentially misleading because the uniramous antenniform appendages of *Yunnanocephalus* and other trilobites were innervated from the deutocerebrum, as in the trilobitomorph *Xandarella* (Strausfeld et al., 2026a). The antennular nerve roots are also visible in *Yunnanocephalus* (Fig. 4A, B). Hence, these cephalic appendages are appropriately referred to as antennules, the uniramous appendages of the deutocerebrum as defined by Boxshall (2004).

### A brain hiding in plain sight

The initial question that persists is why the trilobite brain has remained so elusive. Skeletal features of trilobites obscure possible alignments with mandibulates. In some early Cambrian Olenelloidea, for example, the paired glabellar lobes are not reliable indications of segmentation due to evolved loss or effacement leaving a smooth glabellar surface (Webster, 2009). This does not pertain to *Yunnanocephalus* nor to specimens of the Ordovician *Triarthrus eatoni*, whose renowned pyritization provides exceptional distinction of exoskeletal features. Observations of *Triarthrus* (Park and Kihm, 2017) have proposed that its anterior glabellar lobe (LA) and the following lobes (L3-L0) denote five true segments of the ‘head,’ where LA is associated with the lateral eyes and L3 with the ‘antennae.’ However, that view requires adjustment because the trilobite cerebrum is not segmental. It occupies the space beneath the LA lobe, which likewise does not represent a segment. The cerebrum’s protocerebral domain connects to the lateral eyes; its deutocerebral domain, also beneath the LA lobe, gives rise to the antennular nerves. Applying this organization to *Triarthrus*, its L3 lobe corresponds to the mandibulate T1 segment (tritocerebrum), which is the first true segment providing the first pair of biramous appendages that are not antennae. The three following lobes, L2, L1, and L0, corresponding to mandibulate segments T2, T3, and T4, provide three more identical pairs of biramous appendages, thus resolving four homonomous post-antennular appendages within the cephalon, as recently demonstrated by Hou and Hopkins (2024).

Comparison of the present specimen with other well-described morphologies permits reconstruction of the cerebrum’s position within the trilobite cephalon. As shown in the schematic lateral view (Fig. 6A), the cerebrum occupies the space beneath the LA lobe and whatever space is available forwards of where the LA lobe merges with the sloped cuticle of the fixigena. As described from unique fossil specimens showing the hypostome contiguous with the crop and stomach (El Albani et al., 2025), there would seem to be very little space for a robust cerebrum. This view is contradicted, however, by crowded organs in certain decapod crustaceans, such as the crayfish *Astacus fluviatilis* where, connected to its visual neuropils, a substantial brain rests between the stomach and the frontal wall of the carapace (Huxley, 1896). A comparable disposition pertains to historical soft X-ray images of the Devonian phacopid trilobite *Chotecops* (*Phacops*) *ferdinandi* (Fig. 6B, C), one of many X-ray images of trilobites attributed to the physicist Wilhelm Stürmer (Stürmer et al., 1980; see Schoenemann et al., 2021). The image (Fig. 6B, C) shows a bilaterally symmetric profile of the cerebrum lying above the hypostome and abutting the rostral stomach, as resolved in *Yunnanocephalus* (Fig. 2B, C). Nerves extending forwards are directed towards billowed volumes of fossilized tissue subtending the schizochroal eyes, which typify Phacopida (Schoenemann et al., 2021). A corresponding cerebral disposition has also been identified for the proto- and deutocerebrum of *Anacheirurus adserai* from the Lower Ordovician Fezouata Shale (Losso et al., 2026b). In *Yunnanocephalus* the elongation of the stomach into the narrow dorsal gut occurs immediately behind the location of the LA lobe, hence posterior to the deutocerebrum, whose paired nerve cords would have to extend downwards (as they do in the crayfish) to continue as a ventral arrangement of segmental ganglia. In the compressed *Yunnanocephalus* part (Fig. 2A), the dorsal gut and other tissues obscure the ventral T1 and segmental ganglia further caudally. In *Anacheirurus*, the T1 (‘tritocerebral’) ganglion is visible, clearly preserved as caudal to the esophageal foramen, linked by paired connections to the fossilized rostral deutocerebral outline. Further true segments of the ventral nerve cord are inferred but are deeper in the matrix (see Losso et al., 2026b).

### Trilobite repertoires

Even though the brains of trilobites were generally as modest as the three examples discussed here, they were obviously adapted to meet the demands of their ecology. But what were they? After all, their artiopodan cousins went extinct by the end of the Ordovician, whereas Trilobita persisted for almost another two hundred million years, until the end of the Permian. These considerations return to earlier comments regarding whether the diversity of external traits of the highly conserved trilobite ground pattern would imply different types and distributions of sensory attributes and hence diversification of brain organization across trilobite species. As in contemporary eucrustaceans much of the thoracic and abdominal motor actions would have been governed by local ganglionic circuits (Harzsch, 2003; Zhang et al., 2014), whereas the cerebrum regulates the selection, instigation, and coordination of motor actions for adaptive behaviors, including memory-guided activity. But identifying reasonable proxies amongst today’s marine mandibulates is almost impossible: unlike trilobites, pancrustaceans have evolved a vast range of divergent appendicular specializations and hence elaborate sensory inputs pertaining to the body that are relayed to the cerebrum (Tsubouchi et al., 2017). The remarkable computational capabilities of genetically defined circuits in the minute *Drosophila* brain (e.g., Bates et al., 2026), which is smaller but more derived than the brains of most other hexapod lineages (Ito et al., 2014), suggests that the volume of a trilobite cerebrum was generous, sufficient to orchestrate ecologically appropriate actions. But was their behavioral repertoire as conservative as was their appendicular ground pattern, which remained largely homonomous throughout their evolutionary history? Even the isopods, their marine proxies, possess tagma-related appendicular adaptations for swimming (Hessler, 1993) thus questioning the efficacy of the trilobite legs in a neritic habitat, even though free-swimming predation is attributed to some species (Fortey, 2004). Caution is advised when surmising the attributes of fossil or even extant euarthropod brains. The versatility of thoracic-abdominal appendages is linked to elaborate behavioral repertoires (Vetter and Caldwell, 2015; Patel and Cronin, 2020) and to elaborate central processors (Power, 1943). These typify extant winged insects and free-swimming stomatopods (Thoen et al, 2014). Such computational complexity is rare and remarkable. Their counterparts in euarthropods such as chilopods, are simple layered centers associated with motor actions for limited behavioral repertoires that likely also characterized. basal mandibulates represented by myriapod, leptostracan and archaeognathan pancrustaceans (Loesel et al., 2002; Kenning et al., 2013; Strausfeld et al., 2026b).

## METHODS

### Photomicroscopy

For light microscopy of fossil material, digital images were taken using a 231 Nikon D3X attached to a Leica M205C photomicroscope (Leica Microsystems; Wetzlar, 232 Germany). Images were transferred to Adobe Photoshop CS5 (Adobe Systems; San Jose, CA) and 233 processed using the Photoshop camera raw filter plug-in to adjust sharpness, luminance, texture. Unless otherwise stated, colors were untouched and are those typical of Chengjiang fossils. For ultraviolet illumination (Fig. 2C,D), the fossil’s part was photographed using appropriate filter blocks to evoke intense green fluorescence. Flexible fiberglass light guides were used to combine UV fluorescence and white-light illumination.

### Interpretive drawings and tracings

Drawings were made using Adobe Illustrator for which layered tracings and reconstruction were generated using photographic images as guides.

## Data Availability

The specimen used or referred to here is housed in the University of Arizona Department of Neuroscience, under the curatorship of N. J. Strausfeld. Data reported in this study are included in the article. Any additional information regarding the data reported in this article is available from N.J.S. upon reasonable request.

## Code Availability

This article does not contain original code.

## AUTHOR CONTRIBUTIONS STATEMENT

N.J.S. and F.H. originated the project; N.J.S examined and photographed the specimen of the Lower Cambrian redlichiid trilobite *Yunnanocephalus yunnanensis*. F.H. ascribed published gene expression data to the described panarthropods. N.J.S. and F.H. wrote the manuscript.

## ACKNOWLEDGEMENTS

We thank Professor Michael Montenari, Keele University, Departments of Earth Sciences and Geography, for inviting us to contribute to the second of two issues of ‘Stratigraphy and Timescales’ dedicated to Richard Fortey. Our thanks go to Dr. Camilla Strausfeld for her advice and critical reading of this work. This work has benefitted from the images of developmental stages of *Yunnanocephalus* generously provided by Professor Xingliang Zhang, Director of the Shaanxi Key Laboratory of Early Life and Environment, Department of Geology, Northwest University, China. Likewise, we thank Professor Brigitte Schoenemann for advice regarding the X-ray image of the Devonian trilobite (Fig. 6B) here modified from Fig. 1g in Schoenemann et al., 2021. We also acknowledge the essential importance of Mr. R. Kihm and Professor J. St John’s 2007 translation of, and observations pertaining to, Walch’s 1771 opus, thereby providing present day readers a fascinating view of an enlightened consideration of what trilobites might have been.

